# Pneumococcal attachment to epithelial cells is enhanced by the secreted peptide VP1 via its control of hyaluronic acid processing

**DOI:** 10.1101/788430

**Authors:** Rolando A. Cuevas, Elnaz Ebrahimi, Ozcan Gazioglu, Hasan Yesilkaya, N. Luisa Hiller

## Abstract

The Gram-positive bacterium *Streptococcus pneumoniae* (pneumococcus) is an important human pathogen. It can either asymptomatically colonize the nasopharynx or spread to other tissues to cause mild to severe diseases. Nasopharyngeal colonization is a prerequisite for all pneumococcal diseases. We describe a molecular pathway utilized by pneumococcus to adhere to host cells and promote colonization. We demonstrate that the secreted peptide VP1 enhances pneumococcal attachment to epithelial cells. Transcriptional studies reveal that VP1 triggers the expression of operons involved in the transport and metabolism of hyaluronic acid (HA), a glycosaminoglycan present in the host extracellular matrix. Genetic experiments in the pneumococcus reveal that HA processing locus (HAL) promotes attachment. Further, overexpression of HAL genes in the Δ*vp1* background, reveal that the influence of VP1 on attachment is mediated via its effect on HA. In addition, VP1 also enhances degradation of the HA polymer, in a process that depends on the HAL genes. siRNA experiments to knockdown host HA synthesis support this conclusion. In these knockdown cells, attachment of wild-type pneumococci is decreased, and VP1 and HAL genes no longer contribute to the attachment. Finally, experiments in a murine model of colonization reveal that VP1 and HAL genes are significant contributors to colonization. Our working model, which combines our previous and current work, is that changes in nutrient availability that influence CodY and Rgg144 lead to changes in the levels of VP1. In turn, VP1 controls the expression of a genomic region involved in the transport and metabolism of HA, and these HAL genes promote adherence in an HA-dependent manner. VP1 is encoded by a core gene, which is highly induced *in vivo* and is a major contributor to host adhesion, biofilm development, colonization, and virulence. In conclusion, the VP1 peptide plays a central role in a pathway that connects nutrient availability, population-level signaling, adhesion, biofilm formation, colonization, and virulence.

**AUTHOR SUMMARY:** *Streptococcus pneumoniae* (the pneumococcus) is a major human pathogen. This bacterium asymptomatically colonizes the human upper respiratory tract from where it can disseminate to other tissues causing mild to severe disease. Colonization is a prerequisite for dissemination and disease, such that the molecules that control colonization are high-value candidates for therapeutic interventions. Pneumococcal colonization is a population-level response, which requires attachment to host cells and biofilm development. VP1 is a signaling peptide, highly induced in the presence of host cells and *in vivo*, promotes biofilm development, and serves as a potent virulence determinant. In this study, we build on the molecular mechanism of VP1 function to reveal novel bacterial and host molecules that enhance adherence and colonization. Our findings suggest that host hyaluronic acid serves as an anchor for pneumococcal cells, and that genes involved in the transport and metabolism of HA promote adherence. These genes are triggered by VP1, which in turn, is controlled by regulators that respond to nutrient status of the host. Finally, our results are strongly supported by studies in a murine model of colonization. We propose that VP1 serves as a marker for colonization and a target for drug design.

## INTRODUCTION

The Gram-positive bacterium *Streptococcus pneumoniae* (also known as the pneumococcus) is an important human pathogen. A recent global study on lower respiratory infections determined that the pneumococcus contributed to morbidity more than all other etiologies combined: it was responsible for an estimated 1.18 million deaths (1, 2). This pathogen can either asymptomatically colonize the nasopharynx or spread to other tissues to cause mild to severe diseases. It can spread to the middle ear and sinus, leading to otitis media or sinus infections, and the lungs causing pneumonia (3). It can also disseminate into the bloodstream, brain or heart causing sepsis, meningitis, or heart disease, respectively (4, 5). It is well established that nasopharyngeal colonization is a prerequisite for all pneumococcal diseases. In this study, we explore the mechanisms of colonization and virulence.

Pneumococcus secretes many small peptides, which influence colonization, virulence, and adaptation via effects on competence, intra-species competition, and biofilm development (6–18). The **V**irulence **P**eptide 1 (VP1) is a member of a family of peptides with a conserved N-terminal sequence characterized by a double glycine motif, which directs its export into the extracellular milieu via ABC transporters (7, 19). The *vp1* gene is widely distributed across pneumococcal strains, as well as encoded in related streptococcal species. The gene encoding VP1 is a virulence determinant and is highly upregulated in the presence of host cells where it promotes biofilm development (7, 20). Pneumococcal biofilms have been documented in the nasopharynx, the middle ear, and the sinus (21–24). This mode of growth enhances pneumococcal dissemination and pathogenesis (25, 26). Further, biofilms promote pneumococcal colonization in multiple ways: they facilitate immune evasion, increase resistance to antimicrobials, and provide a platform for DNA acquisition (27–29,29–32). In summary, VP1 is a secreted peptide, induced in the presence of human cells, that promotes biofilm development and virulence.

Glycosaminoglycans (GAGs) are major components of the extracellular matrix. GAGs are acidic linear polysaccharides with a repeated disaccharide unit. The disaccharides consist of an amino sugar (*N*-acetylglucosamine or *N*-acetylgalactosamine) and a uronic acid (glucuronic acid or iduronic acid) or galactose. These polymers are classified into groups based on the nature of the disaccharides, the mode of glycoside bond, and the sulfation levels (33). The main groups are heparin and heparan sulfates (HS), chondroitin and dermatan sulfate (CS), keratan sulfate, and finally hyaluronic acid (HA) (also known as hyaluronate or hyaluronan) (34). Their molecular weights range from 15 to over 100 kDa, and their synthesis takes place in the Golgi apparatus, with the exception of HA, which is synthesized at the membrane (35). GAGs participate in numerous biological processes of the host and the bacteria. In the host, they influence cell adhesion and growth, cell proliferation and differentiation, and tissue formation (for comprehensive reviews, please see Iozzo and Schaefer, 2015; Pomin and Mulloy, 2018; Taylor and Gallo, 2006). In the context of microbial infections, they serve as sources of nutrients and modulators of the immune response (38–41). In addition, many bacteria utilize GAGs to adhere to host cells (42). This includes the pneumococcus, where treatment of lung cells to decrease HS and CS levels reduces pneumococcal adherence (43). In this manner, GAGs are critical components of the bacterial environment during biofilm formation and host interactions.

HA consists of disaccharides of N-acetyl-D-glucosamine (GlcNAc) and D-glucuronic acid (GlcUA). This GAG is present on the apical surface of human bronchial epithelial cells and in the airway mucosa (44, 45). It is also abundant in the synovial fluid, skin, umbilical cord, and vitreous body and is the only non-sulfated glycosaminoglycan in the lung (46–48). This unbranched polysaccharide provides mechanical support, activates immunity, and modulates cell proliferation, migration, and intracellular signaling within the host (49–51). Many streptococcal genomes encode multiple operons for HA transport and metabolism. In the pneumococcus, it has been shown that HA processing and internalization is carried out by a unique set of enzymes including a hyaluronidase, a transporter system, and a hydrolase (52, 53). The current model suggests that HA is digested into disaccharides by a hyaluronidase (*hysA* or *hylA*), predicted to be covalently linked to the cell wall via an LPxTG motif. The disaccharides are imported into the bacteria and phosphorylated via a phosphoenolpyruvate-dependent phosphotransferase system (PTS) system. The PTS system is composed of general and substrate-specific components. Enzyme I (EI) and the histidine-containing phosphocarrier protein (HPr) are cytoplasmic subunits, shared by multiple PTS systems. In contrast, Enzyme II (EII) provides specificity to the PTS via substrate recognition. EII has four enzymes: EIIA and EIIB are cytosolic, and EIIC and EIID are the transmembrane components and form a pore. During internalization via the PTS, the HA disaccharides are phosphorylated. These disaccharides are processed by an unsaturated glucuronyl hydrolase (*ugl*) into the monosaccharide components. Finally, the GlcUA can be further processed into glyceraldehyde 3-phosphate and pyruvate by a set of four genes (the isomerase *kduL*, the reductase *kduD*, the kinase *kdgK*, and the aldolase *kdgA*). Operons for the hydrolase, the PTS-EII system, and the GlcUA metabolism enzymes are all adjacent in the genome, and under the control of RegR, a negative regulator located immediately downstream (54). Note that the unsaturated glucuronyl hydrolase not only cleaves HA into GlcNAc and GlcUA, it can also cleave chondroitin sulfate into their component monosaccharides (53, 55). The HA pathway has been shown to influence pneumococcal biology. Specifically, *hysA* and the genes in the PTS-EII-encoding operon (EIIB, EIIC, EIID, and *ugl*), are required for pneumococcal growth on human HA, and contribute to pneumococcal biofilm formation and virulence (54,56,57).

In this study, we show that VP1 promotes pneumococcal adherence via activation of HA transport and metabolism. Our experiments in a murine model of pneumococcal carriage demonstrate that both VP1 and HA processing promote colonization. We propose a model where the nutritional status of the host induces expression of the pneumococcal secreted peptide VP1, via CodY and Rgg144 (7,18,58). Signaling by VP1 triggers HA metabolism in the bacteria, which in turn, regulates attachment to host cells and colonization in the upper airways.

## RESULTS

### *vp1* enhances pneumococcal attachment to epithelial cells

We have previously shown that *vp1* plays a role in biofilm development (7). Thus, we investigated whether *vp1* influences attachment to epithelial cells, an early step in biofilm formation. For these studies, we utilized strain PN4595-T23, a representative of the drug-resistant and pandemic PMEN1 pneumococcal lineage (59–62). To show that the results extend beyond one lineage, we also employed TIGR4, a clinical isolate frequently used as a model strain (63). For host cells, we utilized the human lung epithelial cells line A549. These cells have been extensively used to study pneumococcal behavior (20,64,65). Early works demonstrated that A549 cells display features of type II alveolar pneumocytes: they are capable of surfactant production (66–68), they produce cell surface-associated MUC1 and secrete MUC5AC mucin in air-liquid cultures (69, 70), and they produce HA. Further, our experiments on adhesion are extended to NIH-3T3 fibroblasts as they express HA and chinchilla middle ear epithelial cells (CMEE) as these primary cells were used in the original characterization of VP1 function (7, 71).

We established that the wild-type strain displays strong attachment to A549 cells after one hour of exposure (Fig S1A). Thus, we used this time point to compare adhesion between wild-type, *vp1* deletion mutant (Δ*vp1*) and *vp1* complement strains (Δ*vp1:vp1*) by using spot plating assays and enumerating the total bound bacteria on TSA plates. The Δ*vp1* displays a reduction in attachment compared to the wild-type cells. Furthermore, the wild-type phenotype is rescued in a *vp1* complement strain (Δ*vp1:vp1*) (Fig 1A). These data suggest that VP1 promotes attachment to the lung epithelia. This role for the *vp1* gene product is not specific to A549 cells, as *vp1* also promotes adherence to chinchilla middle ear epithelial cells (CMEE) and NIH-3T3 fibroblasts (Figs S1B and S1C). Similarly, these results extend beyond the PMEN1 lineage, deletion of the operon encoding *vp1* (*vpo*) in a TIGR4 background also displays decreased attachment relative to the wild-type strain and this phenotype is restored to wild-type levels in a *vpo* complement strain (Fig S1D). Adherent bacteria were visualized by fluorescence microscopy of A549 cells inoculated with PMEN1 cells (Fig 1B). The confocal orthogonal view shows localization at the surface of A549 cells (**Fig S1E**). Finally, the difference in attachment is not the result of variation in growth rate between strains, as growth in DMEM media is the same for wild-type, Δ*vp1,* and Δ*vp1:vp1* strains, as previously reported (7). We conclude that *vp1* promotes pneumococcal attachment to mammalian epithelial cells.

**Fig 1.**
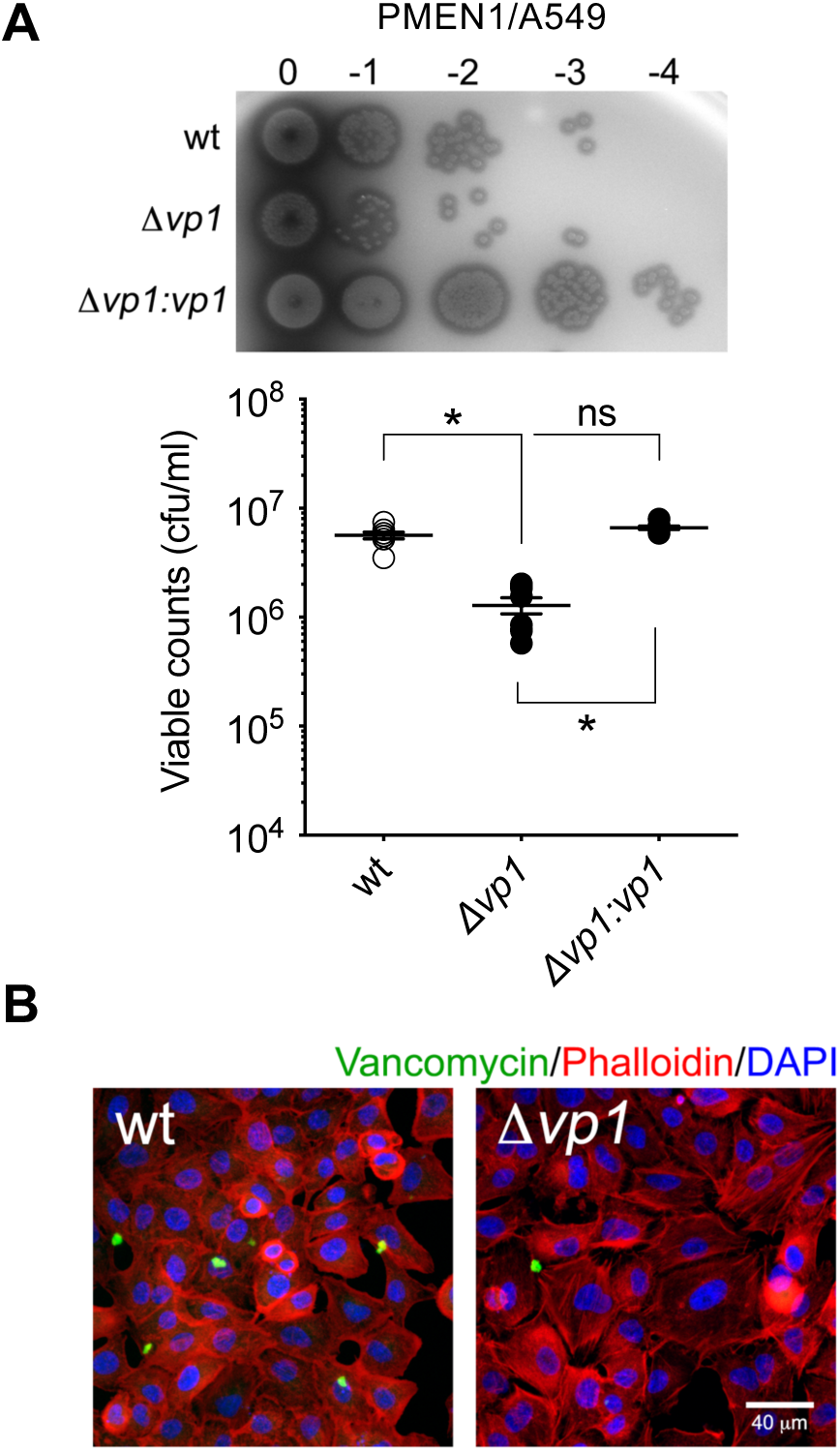
*vp1* enhances pneumococcal attachment to epithelial cells. (A) A549 lung epithelial cells were exposed to the wild-type PMEN1 strain PN4595-T23, and isogenic *vp1* mutant and *vp1* complemented strains (Δ*vp1* and the Δ*vp1:vp1,* respectively) for 1h at 37 °C. After washing the unbound bacteria, the remaining cell-bound bacteria were recovered and quantified on TSA plates. Visualization by a spot assay performed with 5 μl of ten-fold dilutions (top panel). Quantification of the total number of bacteria bound to epithelial cells based on counts from TSA plates (bottom panel). (B) Representative image of wild-type and the Δ*vp1* strains bound to A549 cells. Bacterial cells were stained with BODYPI-FL vancomycin (green), actin was visualized with TRITC-phalloidin (red), and bacterial and host DNA were visualized with DAPI (blue). Data on A represents the mean ± S.E.M of at least six independent experiments. Statistical significance was calculated using unpaired one-way ANOVA analysis with Bonferroni correction, and multiple comparison between samples were performed. * *p* < 0.0001, ns = not significant with *p* = 0.080.

### The *vp1* product positively regulates expression of HA transport and metabolism genes

The VP1 pro-peptide possesses a double glycine motif common to secreted peptides that bind surface-bound histidine kinase receptors from pneumococcal two-component signal transduction system (72). Further, synthetic VP1 binds to the pneumococcal surface (7). Thus, we hypothesized that VP1 signals to producing and neighboring cells and induces expression of genes that participate in the attachment.

To test this hypothesis, we compared the genome-wide expression of the wild-type PN4595-T23 strain and the isogenic Δ*vp1* mutant using the pangenome pneumococcal array (13) (Fig 2A) Strains were grown in chemically defined media (CDM) supplemented with glucose (CDM-Glu), as *vp1* expression is induced in this media relative to rich media (7). In support of the hypothesis, twenty-eight genes displayed a 3-fold difference in expression between the wild-type strain and the Δ*vp1* strain (Fig 2A and S1 Table). Many of these genes are organized into three genomic regions (Fig 2B). Microarray results were validated by qRT-PCR for a subset of genes (S2 Table). Taken together, we observed a significant number of genes with transcription levels that respond to *vp1*.

**Fig 2.**
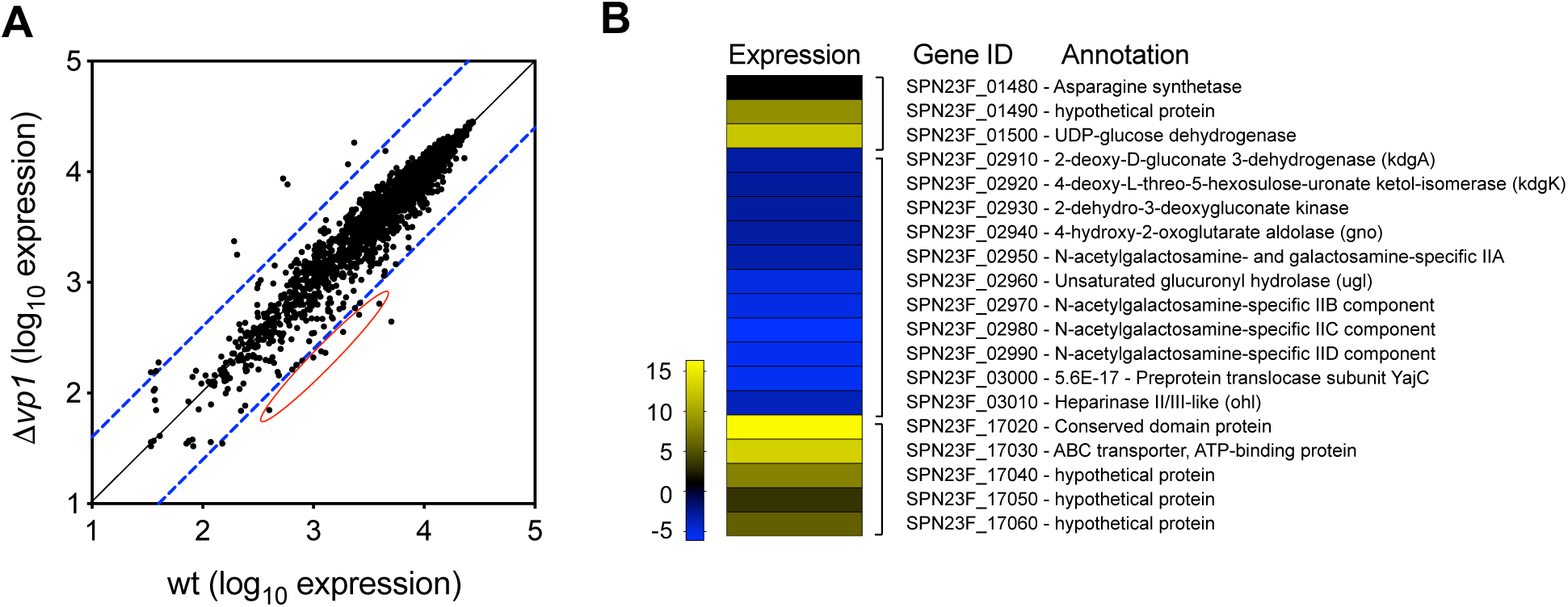
The *vp1* product positively regulates the expression of an operon critical to processing of hyaluronic acid. (A) Microarray transcriptome analysis comparing the expression profiles of the wild-type PMEN1 strain PN4595-T23 and *vp1* derivative strain (Δ*vp1*) on CDM supplemented with glucose. Data was analyzed for statistical significance using CyberT as per Student’s T-test. The dashed blue lines represent 4-fold up- and down-regulation, respectively. The red circle highlights genes associated with hyaluronic acid metabolism. (B) Selected genomic regions with 3-fold difference in expression between the wild-type and the Δ*vp1* strains. Targets were sorted by their genomic position on the genome. Operons are indicated by brackets. Color code illustrates expression ratio between the Δ*vp1* strain relative to wild-type strain. Gene IDs correspond to those in PMEN1 reference strain ATCC700669 (GenBank FM211187), and the predicted functions based on Pfam. Full set presented in S1 Table.

Our analysis shows that two neighboring operons display higher expression in the wild-type strain relative to the Δ*vp1;* these operons have been previously implicated in the processing of HA (53). Together, these two operons encode eleven genes that displayed an average 4.28-fold decrease in expression in the Δ*vp1* relative to the wild-type strain (Figs 2B and 3A). One operon is on the positive strand and encodes seven genes. In this study, for simplicity, we refer to this as the PTS-HAL operon as it encodes the four EIIABCD enzymes that form the PTS importer (Fig 3A). This importer is required for phosphorylation and internalization of HA (EII genes are: SPN23F_02950, and SPN23F_02970 to SPN23F_02990). The operon also contains the *ugl* gene that encodes an unsaturated glucuronyl hydrolase (SPN23F_02960), the preprotein translocase YajC that is a subunit of the bacterial holo-translocon (SPN23F_03000) (73), and finally a predicted heparinase (SPN23F_3010). Our analysis with Phyre2 (74), indicates that the heparinase is a glycosidase that belongs to the superfamily of heparinases type II/III; member of this family bind and degrade heparin and heparin sulfate (75). It is noteworthy, that 60 base-pairs downstream of the end of *hep*, is the negative regulator RegR (SPN23F_3020) (76). Analyses of the region upstream of RegR with the promoter identification programs BDPG and BPROM suggests the presence of an active promoter with the −35 within the *hep* sequence (77, 78). The second operon regulated by VP1 is immediately upstream of the PTS-HAL operon on the negative strand, it encodes four enzymes predicted to metabolize glucuronic acid into glyceraldehyde-3 phosphate and pyruvate (SPN23F_02910 to SPN23F_02940). These transcriptional findings suggest that *vp1* triggers the expression of genes implicated in the transport and metabolism of HA.

**Fig 3.**
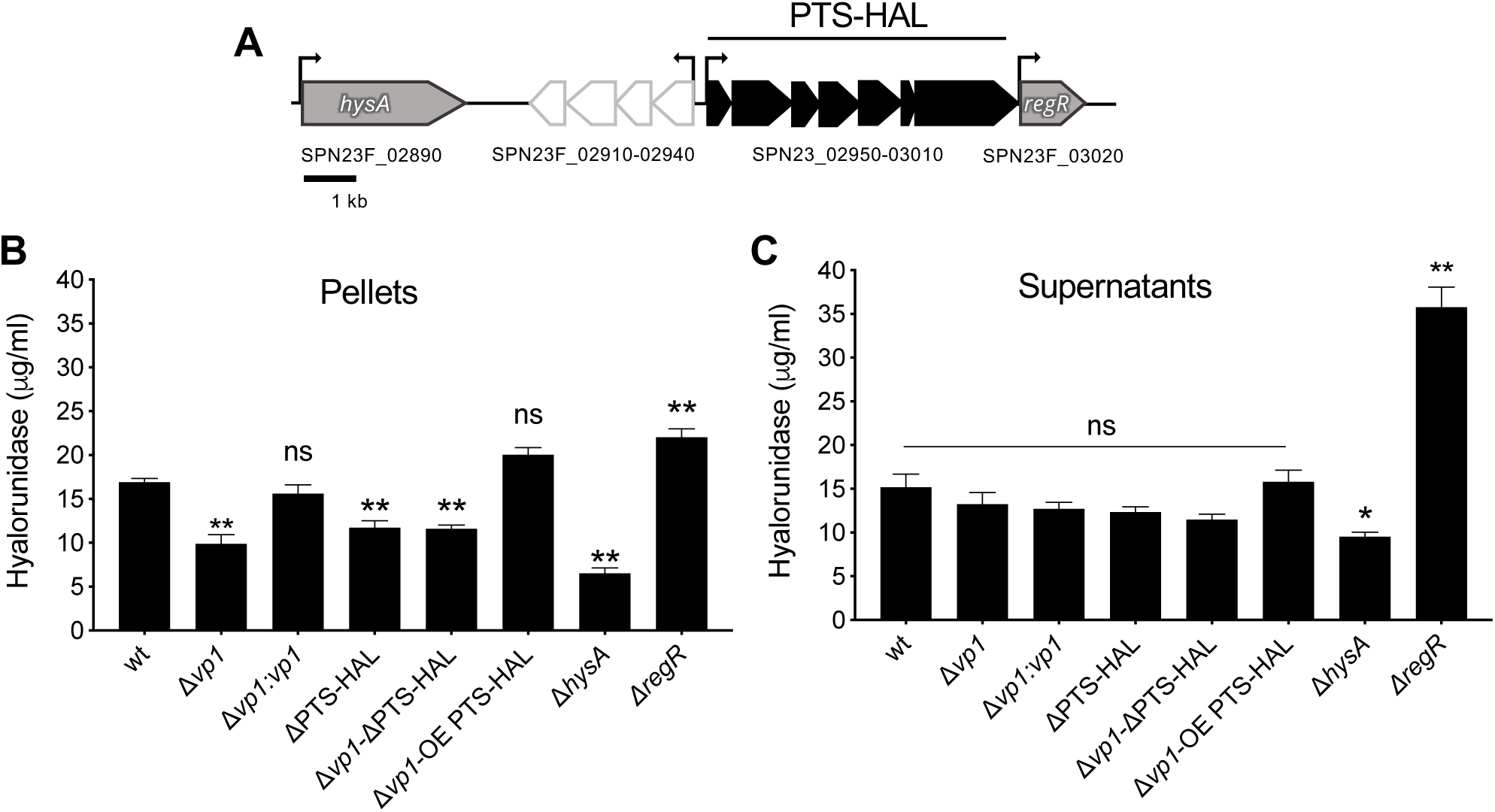
The *vp1* product promotes degradation of the hyaluronan polymer via its controls of the operon encoding the PTS-EII system. (A) Schematic of the genomic region implicated in the metabolism of hyaluronic acid. Arrows correspond to predicted open reading frames. Black and white denote predicted operons. Grey corresponds to genes involved in HA metabolism, but not highly controlled by *vp1*. The IDs refer to gene names in the PMEN1 reference strain ATCC700669. (B and C). Measurements of breakdown of hyaluronan and isogenic deletion mutants for *vp1* (Δ*vp1*), the <u>h</u>yaluronic <u>a</u>cid locus PTS operon (ΔPTS-HAL), *vp1* and the PTS operon (Δ*vp1-*ΔPTS-HAL), *hysA (*Δ*hys)*, and *regR* (Δ*regR*), and for a *vp1* complemented strains (Δ*vp1:vp1*), and a PTS overexpressor in the Δ*vp1* background (Δ*vp1*-OE PTS-HAL). Both the pellets (B) and the supernatants (C) of bacteria grown in liquid cultures were collected and assessed for hyaluronidase activity. Data on B and C represents the mean ± S.E.M of at least six independent experiments. Statistical significance was calculated using unpaired one-way ANOVA analysis with Bonferroni correction, and multiple comparison between samples were performed. * *p* < 0.05, ** *p* < 0.001, ns = not significant relative to the wild-type strain. Note that on B, the complemented Δ*vp1:vp1* rescues the hyaluronidase deficiency of the Δ*vp1* mutant (*p* < 0.0001) and the overexpressor Δ*vp1-*OE PTS-HAL rescues the hyaluronidase activity of the mutants Δ*vp1* (*p* < 0.0001). Further, there is no significant difference between Δ*vp1* and ΔPTS-HAL, between Δ*vp1* and Δ*vp1-*ΔPTS-HAL.

### The *vp1* product promotes degradation of the HA polymer via its control of the HA processing and transport operon

Once we established that VP1 stimulates expression of multiple genes involved in HA metabolism, we tested whether VP1 could influence the processing of the HA polymer. To this end, we measured the hyaluronidase activity in pellets and supernatants of the PN4595-T23 strain carrying deletions of *vp1,* PTS-HAL, and various genes involved in HA processing (Fig 3B). In this assay, bacterial pellets or supernatants were mixed with the HA polymer, degradation of HA was measured as a change in optical density at OD_400_ and reported as hyaluronidase concentration (76). As controls, we employed the Δ*hysA*, as this gene encodes for a hyaluronidase and its deletion should result in reduced hyaluronidase relative to the wild-type strain (53, 79). As predicted, this strain displayed 2.6-fold lower hyaluronidase levels (*p* < 0.0001 relative to the wild-type strain). We also employed a Δ*regR* mutant, as this gene encodes for a repressor of *hysA* and its deletion should lead to increased hyaluronidase activity (54). The Δ*regR* strain exhibited elevated hyaluronidase activity (*p* = 0.0006). The Δ*vp1* strain displayed 1.7-fold lower hyaluronidase compared to the wild-type strain (*p* < 0.0001). This defect was rescued in the Δ*vp1:vp1* (*p* < 0.0001 relative to Δ*vp1* strain). The ΔPTS-HAL mutant and the Δ*vp1-*ΔPTS-HAL double mutant showed a 1.4- and 1.5-fold reduction in hyaluronidase relative to the wild-type strain (*p* < 0.0005). This reduction was similar to the Δ*vp1* strain compared to the wild-type strain. The majority of the hyaluronidase activity was captured within the pellets, suggesting that the majority of the degradation enzymes are associated with the cell and not secreted into the supernatant. These data suggest that *vp1* and the PTS-HAL processing and transport operon promote degradation of HA.

To determine whether the ability to break down HA is conserved beyond the PMEN1 strain PN4595-T23, we performed the same set of experiments in a TIGR4 background. The locus in TIGR4 resembles that of the PMEN1 strain, with the addition of a predicted transposon at the end of the operon (see graphic Fig S2A). The results were similar in that the genes encoded by the *vp1* operon (*vpo*) and HA processing and transport operon (PTS-HAL) contributed to the degradation of the HA polymer. In contrast to PN4595-T23, the majority of the TIGR4 activity was localized to the supernatant (Fig S3C), suggesting differences in the subcellular localization of the hyaluronidase activity between strains.

Finally, to investigate whether VP1 exerts its influence via genes encoded in the HA processing and transport operon, we tested whether overexpression of the PTS-HAL operon in a Δ*vp1* background could rescue the wild-type phenotype in both TIGR4 and PN4595-T23 strains. To this end, using the Δ*vp1* backgrounds, we replaced the native promoter of PTS-HAL with a constitutive promoter (from *amiA* gene) (strain Δ*vp1-*OE PTS-HAL)(80). These strains displayed HA degradation comparable to the isogenic wild-type strains (*p* = 0.21) (Fig 3B and S2C). These data suggest that *vp1* exerts its effect on HA metabolism via control of the gene products encoded in the PTS-HAL operon.

### Pneumococcus attaches to host HA in a process controlled by VP1 and mediated by molecules involved in HA processing

We have demonstrated that the *vp1* product enhances attachment of pneumococci to lung epithelial cells (Fig 1), and promotes transcription and activity of genes in the HA processing and transport operon associated with HA metabolism (Figs 2 and 3). Thus, we hypothesized that *vp1* influences attachment via its effect on the PTS-HAL operon. To test this hypothesis, we performed attachment assays in a strain with a deletion in the PTS-HAL operon (ΔPTS-HAL); it displayed decreased attachment relative to the wild-type strain (Fig 4A). Moreover, the attachment defect observed in the Δ*vp1* mutant was rescued to wild-type levels in the Δ*vp1*-OE PTS-HAL strain where the genes in the PTS-HAL operon are expressed from a constitutive promoter. These data strongly suggest that the products of *vp1* and PTS-HAL promote adhesion and that the effect of *vp1* is exerted via its influence on PTS-HAL. Further, attachment levels for the double mutant Δ*vp1*ΔPTS-HAL resembled that of the ΔPTS-HAL strain consistent with these gene products acting within the same pathway (Figs S3A and S3B). The Δ*regR* strain, with increased levels of expression for genes encoded in the PTS-HAL operon, resembled wild-type levels. The Δ*hysA* strain displayed similar attachment to the wild-type strain, suggesting that genes in PTS-HAL are not promoting attachment via indirect effects on HysA (Fig S3A). The attachment of the wild-type and mutant strains were also visualized by confocal microscopy, and representative images are displayed in Fig 4B. Together, these data demonstrate that the *vp1* promotes attachment via its control of the genes in the PTS-HAL operon, responsible for processing and transport of HA.

**Fig 4.**
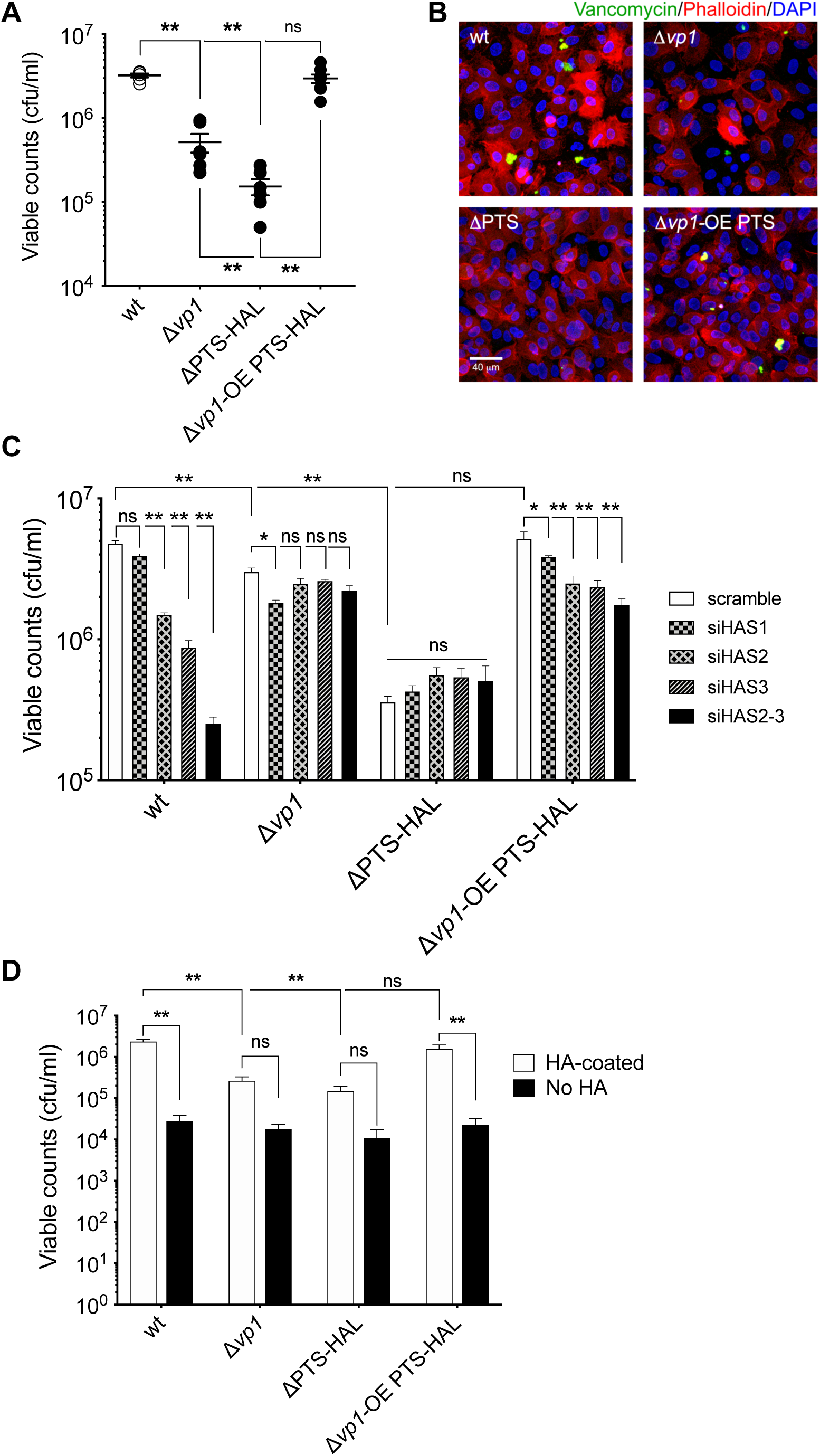
Pneumococcus attaches to host HA in a process controlled by VP1 and mediated by molecules involved in HA processing. (A) The wild-type PN4595-T23, Δ*vp1*, ΔPTS-HAL, and Δ*vp1-*OE PTS-HAL strains were tested for attachment to epithelial A549 cells. Cell bound bacteria were enumerated by plating on TSA plates. (B) Representative image of strains attached to epithelial A549 cells. Bacterial cells were stained with BODYPI-FL vancomycin (green), actin was visualized with TRITC-phalloidin (red), and DNA was visualized with DAPI (blue). (C) Adherence measurements on A549 cells. *HAS1-, HAS2-,* and *HAS3-*knockdown, and the double-knockdown of *HAS2* and *HAS3* (HAS2-3) cells were exposed to wild-type and its derivative Δ*vp1*, ΔPTS-HAL, and Δ*vp1-* OE PTS-HAL strains. siRNA treatment was administered 48 h before addition of bacteria, and adherent bacteria were enumerated by plating on TSA plates. Bound bacteria were enumerated by plating cells on TSA plates. (D) The wild-type PN4595-T23 and its derivative Δ*vp1*, ΔPTS-HAL, and Δ*vp1-*OE PTS-HAL strains were assessed for attachment to hyaluronic acid-coated surfaces in FBS-supplemented DMEM media. The total cell-bound bacteria were recovered and quantified by growth on TSA plates. Data on A, C, and D represents the mean ± S.E.M of at least four independent experiments. Statistical significance was calculated using unpaired one-way ANOVA (A and D) or two-ways ANOVA (C) analysis with Bonferroni correction, and multiple comparison between samples were performed. * *p* < 0.05, **** *p* < 0.0001, ns = not significant. Note that the overexpressor Δ*vp1-*OE PTS-HAL rescues the attachment deficiency of the mutant Δ*vp1* (*p* < 0.0001) in D.

HA production in mammalian cells is controlled by at least three different, membrane-anchored hyaluronan synthases: *HAS1*, *HAS2*, and *HAS3* (35). Previous transcriptomic studies have reported that A549 cells exclusively express *HAS2* and *HAS3*, while *HAS1* remains undetectable (81). Our qPCR analysis corroborates these findings (Fig S3C). To evaluate whether the HA produced by the A549 cells is implicated in pneumococcal attachment, we employed siRNA to knockdown the expression of *HAS2* and *HAS3* (independently and together) on A549 cells, and 48 h later, assessed the attachment strength of the wild-type strain. As a control for treatment, we used a scrambled construct for siRNA experiments, as well as *HAS1* because it has the low expression levels in A549 (Fig S3C and as described by Chow et al., 2010). At the 48h time point, the siRNA treatment reduced levels of HAS2 and HAS3 to below 20% when compared to the scrambled control (Fig S3D). Our attachment assays revealed a significant reduction in pneumococcal attachment to host cells where HAS2 or HAS3 were knockdown, with maximum reduction in the double knockdown cells (Fig 4C). These data suggest that pneumococcus binding is enhanced in the presence of host HA.

If products of the *vp1* gene and the PTS-HAL operon exert their influence on attachment via HA, a dramatic decrease in host HA levels should reduce the effect of these pneumococcal molecules on host attachment. Consistently, the Δ*vp1* strain displayed similar levels of attachment on host cells independent of the levels of HAS2 HAS3. Further, the ΔPTS-HAL strain also displayed the same levels of attachment in host cells regardless of the levels of HAS2 and HAS3 (Fig 4C). Moreover, attachment of the ΔPTS-HAL strain resembled levels of a wild-type strain on cells with reduced HAS2 and HAS3, that is, substantially lower than attachment to wild type cells. Finally, a strain overexpressing the PTS-HAL operon in the Δ*vp1* background attached better in host cells that express HA. Thus, pneumococcus binding is enhanced by host HA, in a process that depends on the products of *vp1* and PTS-HAL.

The influence of HA in promoting pneumococcal binding is further corroborated by an experiment with exogenous HA. In one experiment, the addition of exogenous HA diminishes binding to A549, consistent with bacteria binding to detached HA (Fig S3E). In a second experiment, we coated 6-well plates with HA from *S. equis* (Fig S4A). A spot assay revealed that pneumococcal attachment was substantially higher on a surface coated with HA, relative to those without HA (Fig S4B). As observed on cells, the levels of attachment for Δ*vp1* or ΔPTS-HAL strains were lower than that of the wild-type strain in HA-coated surfaces, and overexpression of the PTS-HAL operon in the Δ*vp1* restored attachment to wild-type levels (Fig 4D). Moreover, these genetic changes had no significant effect on binding on surfaces that were not treated with HA. All together, these data demonstrate that the products of genes involved in the HA processing and transport promote the binding of pneumococcus to host HA in a process regulated by VP1.

### VP1 and the HA processing and transport operon promote pneumococcal colonization

To establish the role of VP1 and of the PTS-HAL operon in pneumococcal survival in the nasopharynx, we employed the murine model of colonization. Cohorts of five mice were inoculated with single strains, using an inoculation dose of 20μl, where the wild-type strain colonizes the nasopharynx, but not the lungs (83, 84). The bacterial load was measured two hours and seven days post-inoculation. Enumeration of pneumococci in the nasopharyngeal lavages two hours post-infection did not show any difference in bacterial load among the strains, indicating the accuracy of infection and the consistency in obtaining nasopharyngeal content (Fig 5A). Throughout the course of infection, the number of recovered wild-type pneumococci remained constant (Figs 5A and 5b).

**Fig 5.**
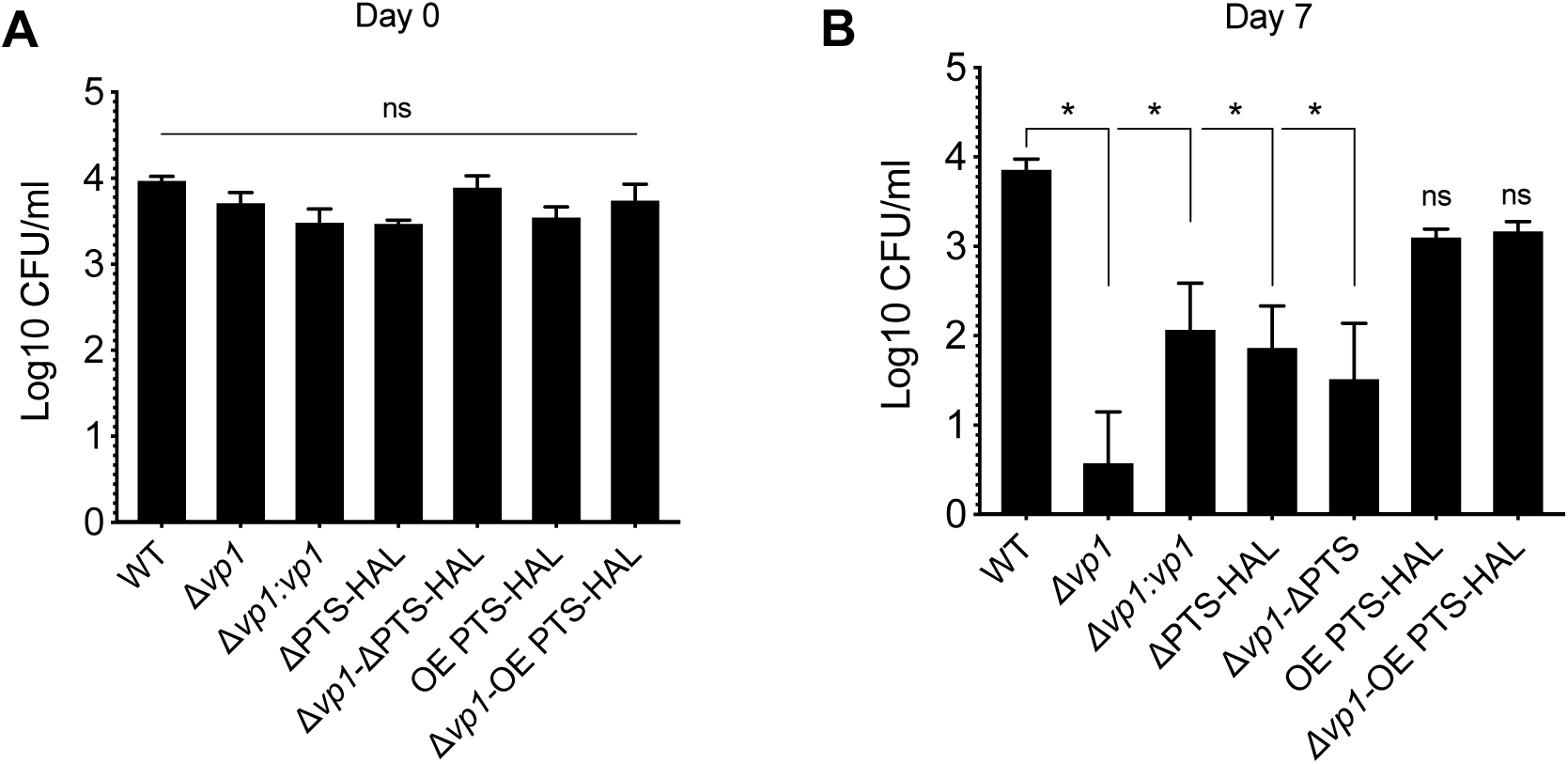
HA processing region is required for pneumococcal colonization. CD1 mice were inoculated intranasally with 20 ml PBS containing approximately 5X10^5^ CFU/mouse. At 2 hours (day 0, A), and 7-days (B) post inoculation, nasopharyngeal contents were collected and pneumococci were enumerated. Each column represents the mean of data from five mice. Error bars show the standard error of the mean. Significant differences in bacterial counts are seen comparing to the wild-type strain using two-way ANOVA and Tukey’s multiple comparisons test. * *p* < 0.0005, ns = not significant. Note that at day 7, the complemented Δ*vp1:vp1* rescues the colonization deficiency of the Δ*vp1* mutant (*p* = 0.003). Similarly, the overexpressor OE PTS-HAL rescues the colonization deficiency of the mutants Δ*vp1* (*p* < 0.0001). The ΔPTS-HAL (*p* = 0.0001) and the double-mutant Δ*vp1-*ΔPTS-HAL (*p* = 0.0001) both are significantly different from the wild-type strain and the colonization deficiency in the Δ*vp1* strain is greater to the ΔPTS strain (p = 0.0176).

Samples recovered seven days post-inoculation revealed significant differences in colony counts across the pneumococcal strains (Fig 5B). Our previous work, in the chinchilla model of pneumococcal disease, established that *vp1* is a virulence determinant based on the reduced mortality and decreased dissemination associated with the *Δvp1* strain relative to the wild-type strain (7). Akin to the chinchilla model, the *Δvp1* strain displayed a dramatic decrease in the murine model, where four out of five animals cleared the infection (*p* < 0.0001). Complementation of *vp1* (Δ*vp1:vp1*) partially rescued the wild-type phenotype. The difference between wild-type and complement (Δ*vp1:vp1*) is likely due to changes in the dose driven by variation of the promoter. These data suggest that VP1 contributes to infection in multiple tissue types, as well as in carriage and disease.

By the seventh day post-inoculation, we also detected a significant reduction in colony counts in the ΔPTS-HAL and the double Δ*vp1-*ΔPTS-HAL mutants compared to the wild-type strain (*p* < 0.0001). Moreover, overexpression of the PTS-HAL operon in the Δ*vp1* background significantly increased the recovered pneumococci compare to Δ*vp1* (*p* < 0.001). Yet, overexpression of PTS-HAL in the wild-type background had no effect (p = 0.542). The observation that the PTS-HAL operon can compensate for the absence of *vp1* is consistent with the model where VP1 acts upstream of HA processing (Figs 5 and 6). These results strongly suggest that VP1 and HA transport and metabolism play a critical role in pneumococcal colonization of the upper airways.

**Fig 6.**
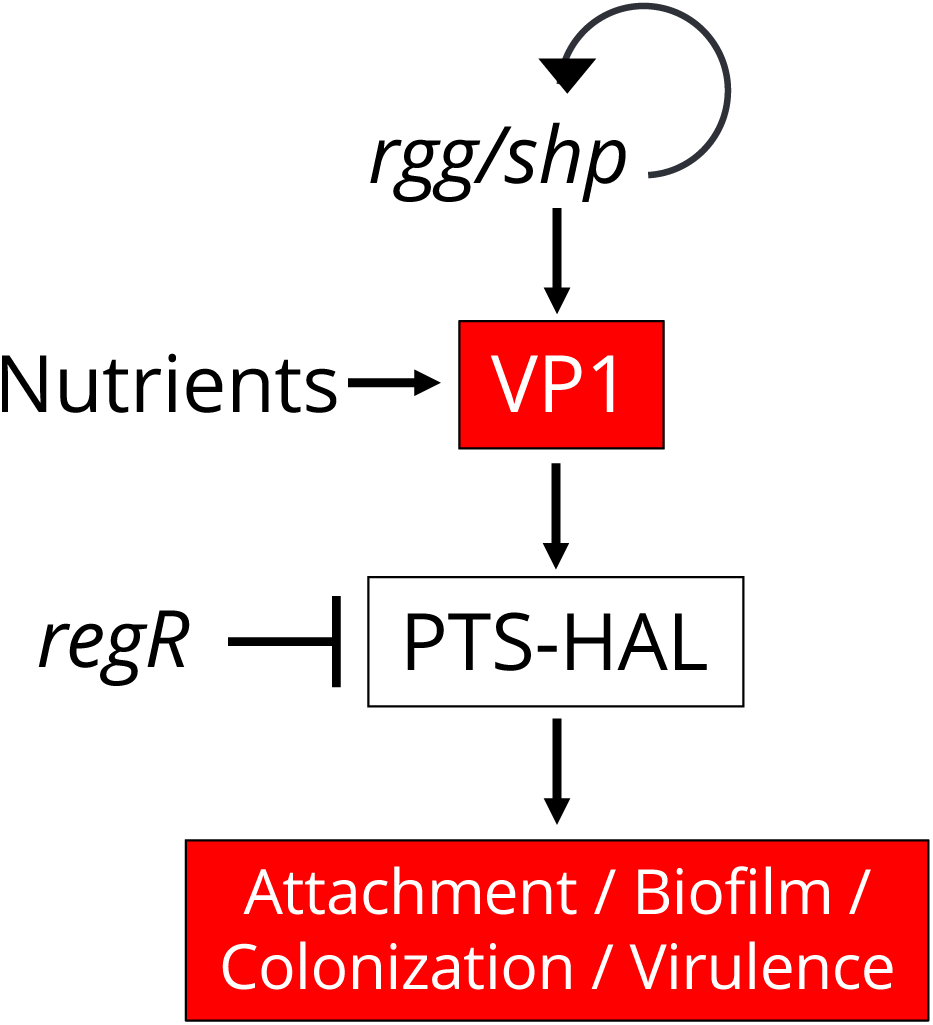
Working model for VP1 regulation and its role in cell-cell communication in *Streptococcus pneumoniae*. The VP1 is processed and secreted into the extracellular milieu. *vp1* is highly upregulated in the presence of host cells and is induced by an upstream transcription factor, Rgg/Shp. In addition, previous work shows that levels of *vp1* respond to nutrients and to the master regulator CodY. VP1 induces changes in the expression of multiple operons, including two implicated in the processing and acquisition of hyaluronic acid. We propose that VP1 promotes attachment via its influence on the products of genes encoded in the PTS-HAL operon.

## DISCUSSION

The binding of pneumococci to epithelial cells of the host mucosal surfaces is a crucial step during colonization and precedes dissemination to other tissues (3,85,86). In this study, we reveal a pneumococcal pathway that promotes adherence via host HA. We previously documented that the signaling peptide *vp1* is dramatically induced during host infection, enhances biofilm formation, and is a potent virulence determinant in a chinchilla model of middle ear infection (7). Here, we build on this work, providing mechanistic details that reveal links across signaling, adherence, and colonization. Specifically, we demonstrate that VP1 controls genes involved in the transport and metabolism of the host glycosaminoglycan, HA. Our *in vitro* studies demonstrate that genes in the PTS-HAL operon enhance attachment of the pneumococcus to epithelial cells. Our *in vivo* studies on a murine model of colonization suggest that VP1 and the operon for HA transport and metabolism play critical roles in nasopharyngeal colonization.

VP1 is part of a regulatory network involved in nutritional sensing. Previous work demonstrated that the master nutritional regulator CodY negatively regulates the level of vp1 transcripts, and positively regulated by the transcriptional regulator Rgg144 (18, 87). The activity of Rgg144 (nomenclature based on gene name in D39, SPD_0144) is regulated by a cognate peptide SHP144 (<u>s</u>hort <u>h</u>ydrophobic <u>p</u>eptide 144), which is secreted from cells and internalized into producing and neighboring cells where it binds to Rgg144 (7, 18). Rgg144 is a negative regulator of the capsule: it directly binds a predicted promoter upstream of the capsular genes, inhibits the expression of capsular genes, and leads to a reduction in the size of the type 2 capsule (18). The bacterium capsule hinders cell adhesion by concealing bacterial receptors (88). In addition, our work on murine models shows that the Rgg144-SHP144 system serves as a colonization factor in the nasopharynx and contributes to pathogenesis in pneumonia (18). These data are consistent with a model where the Rgg144-SHP144 system plays an essential role in colonization by decreasing the capsule and inducing *vp1*, and that together these events promote adhesion to epithelial cells in the nasopharynx and the lungs. Of note, adhesion experiments in this study were performed in CDM supplemented with glucose. In this condition, levels of *rgg144* are low, and levels of *vp1* are high (7, 18). Thus, while Rgg144-SHP144 likely plays a key role in multiple aspects of colonization in our animal model, it is unlikely to be a major contributor *in vitro* experiments presented in this study. We propose that VP1 is a critical component of a complex cellular response to changes in nutritional status.

Our studies in the murine model of colonization reveal a dramatic defect for the *vp1* deletion mutant, where four out of five mice cleared the infection (Fig 5). The strain with a deletion of the HA transport and metabolism operon also displayed a decrease in bacterial load, but not of the same magnitude as the Δ*vp1*. These data suggest that: (1) VP1 mediates attachment via control of the HA transport and metabolism operon, and (2) VP1 plays additional roles in colonization, which are mediated by yet unknown factors.

Many pneumococcal surface molecules serve as adhesins, mediating attachment of the bacteria to host cells (40,89,90). A subset of adhesins binds to extracellular matrix components. Pneumococcal virulence protein A and B (PavA and PavB) (91–93) and enolase (94, 95) bind to fibronectin and plasminogen. Similarly, the plasmin- and fibronectin-binding proteins A and B (PfbA and PfbB) bind to the extracellular matrix and are required for adherence to lung and laryngeal epithelial cells (96, 97). The choline-binding protein A (CbpA or PspC) binds vitronectin (98), and PsrP binds to keratin (99). Pneumococcus also encodes surface-exposed glycosidases, which have been shown to contribute to adhesion not only via modification of the host surface that reveal receptors but also via direct binding (100). The glycosidase NanA (64) and the β-galactosidase BgaA (101), both process GAGs. NanA releases terminal sialic acid residues and BgaA releases terminal galactose residues from glycoconjugates. However, these enzymes promote attachment to host cells in a manner independent of their enzymatic activity. This observation is particularly evident for BgaA, where binding is mediated via a carbohydrate-binding module (102). Thus, the pneumococcal can employ glycosidases in host attachment. In this study, deletion of the surface exposed HysA strongly influenced HA processing (Fig 3), but not attachment (Fig 4). Thus, we deduce that the glycosidase HysA does not mediate the VP1-regulated defect in adhesion.

The data presented in this study strongly suggest that the transport and processing of host HA plays an essential role in adhesion. However, we have not yet established the molecular mechanism responsible for this outcome. Levels of regulators, carbohydrates, and transporters are interconnected in the bacterial cells via highly complex networks. Regulators control gene expression, including levels of transport systems and biosynthesis enzymes. Carbohydrates can function as substrates, and can positively or negatively influence the activity of regulators. Transporters not only control the import of carbohydrates, but the extent and position of phosphorylated residues on the cytoplasmic Hpr in the PTS system also controls CcpA-dependent carbon catabolite repression (103). CcpA influences the levels of almost one-fifth of pneumococcal genes, including ABC transporters and regulators (103, 104). Thus, many molecular mechanisms may explain the connections between transporters and adhesions.

The HA transport and metabolism locus manipulated in this study encodes all the PTS-EII components (EIIA-EIIB, EIIC, and EIID) that specifically mediate HA import. The locus also encodes two additional genes for a putative heparinase and YajC, a component of the machinery associated with the secretion of proteins with N-terminal signal sequences (73), Data presented in Fig 4, where host HA levels are decreased using SiRNA knockdowns, strongly suggest that the PTS and VP1 mediate attachment, via a yet uncharacterized effector, in an HA-dependent manner.

Multiple lines of evidence suggest that GAGs facilitate attachment of multiple bacterial species to host cells. Several groups have explored and advanced the utilization of HA to facilitate attachment of bacteria to abiotic surfaces (49,105,106). A recent report showed that multidrug-resistant *Staphylococcus aureus* and *Pseudomonas aeruginosa* form more robust biofilms on surfaces coated with collagen and HA than on uncoated control surface (42). In addition, levels of GAGs on A549 cells were positively associated with the attachment of several pathogens, including *Staphylococcus aureus*, *Streptococcus pyogenes*, and the pneumococcus (43). Similarly, attachment of multiple bacterial species (both Gram-positive and Gram-negative) was positively associated with GAG levels on corneal epithelial cells HCE-2, well known for expressing a variety of GAGs on their surface (40). In yet another example, HA is the primary attachment site of *M. tuberculosis* in A549 cells (107). While HA has not been directly implicated in the attachment of pneumococcus, previous studies demonstrate that the pneumococcus can utilize HA as a carbon source. In particular, *hysA* and genes in the PTS-EII-encoding operon (EIIB, EIIC, EIID, and *ugl*), are required for pneumococcal growth in media where HA is the only carbohydrate (56). Furthermore, HA supports *in vitro* pneumococcal biofilm formation (57). These studies suggest that multiple bacteria may utilize host HA for attachment or nutrients and that control of each or both of these processes influences colonization and virulence.

In this study, we reveal a novel pathway involved in the attachment of the pneumococcus to host epithelial cells. Our working model is that Rgg144 and CodY monitor nutrient availability and, both of these transcription factors influence levels of the secreted peptide *vp1*. VP1, in turn, controls a genomic region involved in the transport and metabolism of HA, and these genes promote adherence in an HA-dependent manner. VP1 is encoded by a core gene, is highly expressed during infection, and is a significant contributor to both colonization and virulence. Thus, this molecule is a high-value candidate for the development of anti-pneumococcal therapies.

## MATERIALS AND METHODS

### Bacterial strains and culture conditions

Wild-type *S. pneumoniae* strain PN4595-T23 (GenBank ABXO01), was selected as a representative of the PMEN1 lineage (59–62). The clinical isolate TIGR4 (<u>T</u>he Institute for <u>G</u>enomic <u>R</u>esearch, Aaberge et al., 1995), was graciously provided by Dr. Jason Rosch. The construction of the *vp1* deletion mutant (Δ*vp1*) and the *vp1* complemented strain (Δ*vp1:*Δ*vp1*) in the PN4595-T23 background was reported previously (7). Strain PN4595-T23 was modified to also generate the PTS-HAL operon deletion (ΔPTS-HAL), the *hysA* deletion (Δ*hysA*), and the *regR* deletion (Δ*regR*). We generated a double mutant *vp1*-PTS (Δ*vp1*-ΔPTS-HAL) and a PTS-HAL overexpressor in the Δ*vp1* and wild-type backgrounds (Δ*vp1*-OE PTS-HAL and OE PTS-HAL, respectively) (S5 Table). A similar set of strains was constructed in the TIGR4 background (S5 Table). In this set, we generated a deletion mutant strain of the operon encoding *vp1* (*vpo*) and the *vpo* complemented strain (Δ*vpo*::Δ*vpo*). For growth on solid media, strains were streaked onto Trypticase Soy Agar II plates (TSA) containing 5% sheep blood (BD BBL, New Jersey, USA). For growth in liquid culture, colonies from a frozen stock were grown overnight on TSA plates and inoculated into Columbia broth (Remel Microbiology Products, Thermo Fisher Scientific, USA). The media was supplemented with antibiotics as needed. Cultures were incubated at 37**°**C and 5% CO_2_ without shaking. Experiments in chemically defined medium (CDM) were performed utilizing a published recipe (104), and glucose was used at a final concentration of 55mM. Growth in CDM was initiated by growing a pre-culture in Columbia broth for 2-3 hours and then diluted to an absorbance of 0.1 at 600 nm, this culture was then grown in CDM-Glu.

### Strain construction and transformation

To generate the ΔPTS-HAL mutant strain we used site-directed homologous recombination to replace genes SPN23F_02950 to SPN23F_0310 with a kanamycin-resistance gene (*kan*). Likewise, the PTS-HAL operon in the TIGR4 strain (SP_0321 to SP_0327) was replaced with *kan*. The *kan* region was originally amplified from an *rpsL* cassette (80). Gene IDs for PN4595-T23 and TIGR4 are listed in S3 Table. The same strategy was used for the single gene replacements, of *regR* (SPN23F_03020) and *hysA* (SPN23F_02890). To decrease the likelihood of polar effects in all our constructs, the strong artificial transcriptional terminator B1002 was inserted downstream of *kan* (109). Briefly, the transforming DNA was prepared by ligation of the kanamycin gene and promoter, with DNA (one to two kilobases) from the flanking regions upstream and downstream of the PTS-HAL operon. Flanking regions were amplified from the parental strains using either Q5 polymerase or OneTag polymerase (New England Biolabs). The OE PTS-HAL was generated in the Δ*vp1* and wild-type backgrounds by replacing the native PTS operon promoter with an *amiA* promoter and *kan* gene. Assembly of these transforming fragments was achieved by either sticky-end ligation of restriction enzyme-cut PCR products or by Gibson Assembly using NEBuilder HiFi DNA Assembly Cloning Kit. The resulting constructs were transformed into PN4595-T23 or TIGR4. Primers used to generate the constructs are listed in S4 Table. PN4595-T23 and TIGR4 strains were transformed with approximately 1 µg of DNA. Liquid cultures were supplemented with 125 μg ml^−1^ of CSP1 (EMRLSKFFRDFILQRKK, for TIGR4) or CSP2 (EMRISRIILDFLFLRKK, for PN4595-T23), at an absorbance of 0.05 at 600 nm (GenScript, NJ, USA), and incubated at 37°C. After 2 hours, the treated cultures were plated on Columbia agar containing the appropriate concentration of antibiotic for selection, spectinomycin, 100 μg ml^-1^, kanamycin 150 μg ml^-1^). Resistant colonies were cultured in Columbia broth. Sequences were confirmed using DNA sequencing of the PCR amplimers (Genewiz, Inc., USA). All strains generated in this study are listed in S5 Table.

### RNA purification, reverse transcription (RT) and qPCR

Bacterial samples were collected on RNALater (Thermofisher), and pellets were lysed with 1x lysis mix containing 2 mg ml^-1^ of proteinase K, 10 mg ml^-1^ of lysozyme, and 20 μg ml^-1^ mutanolysin in TE buffer (10 mM Tris·Cl, 1 mM EDTA, pH 8.0) for 20 min. Total RNA was isolated using the RNeasy (Zymo Research) following manufacturer instructions. Contaminant DNA was removed by incubating total RNA samples with DNase (2U/μl) at 37°C for at least 15 min and then checked by amplification of *gapdh* (no visible band should be observable in RNA only samples). RT reaction was performed using 1 μg of total RNA using SuperscriptVILO kit for 1 h. Five ng of total cDNA was subjected to real-time PCR using PowerUp SYBR Green Master Mix in the ABI 7300 Real-Time PCR system (Applied Biosystems) according to the manufacturer’s instructions. All qRT-PCR amplification was normalized to pneumococcal *gapdh* and expressed as fold change with respect to the wild-type strain. qPCR primers are listed in S4 Table. Primers were obtained from IDT (Integrated DNA Technologies).

### Spot assays and CFU counts

Bacteria were inoculated from overnight plates into 20 ml of Columbia broth. Liquid cultures were grown statically and monitored at an optical density of 600 nm using Nanodrop 2000c spectrophotometer (Thermo Scientific). Bacterial counts were assessed by streaking 10-fold dilutions on TSA or TSA-antibiotic containing plates. Spot assays were performed with 5 μl of ten-fold dilutions.

### Microarray Analysis

We utilized the Pneumococcal Supragenome Hybridization Array (SpSGH) to compare gene expression between the wild-type PN4595-T23 strain and the Δ*vp1* as described previously (13). Pneumococcal cultures were collected and normalized to an absorbance of 0.3 at 600 nm. RNA extraction, cDNA preparation, and cDNA labeling were performed as described previously (110). Cyber T was used for data analysis (111). Microarray experiments were performed in triplicate. Genes with at least a 3-fold difference between strains and a Bayesian *p*-value < 0.05 were selected for further analysis. Microarray data for selected genes was confirmed by qPCR.

### Preparation of HA-coated surfaces

Hyaluronic acid from *Streptococcus equi* (Sigma-Aldrich) was bound to tissue culture 24-wells plates following a previously described method with modifications (49). Briefly, twenty-four well plates (VWR) were treated with 0.5 ml of concentrated sulfuric acid for 10 min at 60**°**C, washed extensively with distilled water, then incubated for 2 h at 37**°**C with 0.5 ml 5 mg ml^-1^ of hyaluronic acid. Finally, the plates were rinsed with distilled water and allowed to stand for 1h at room temperature. Plates were not dried during the entire procedure. Control plates were treated solely with either sulfuric acid or HA alone. Hyaluronic acid binding was assessed with 0.5 ml of 1% of Alcian Blue 8GX (Sigma-Aldrich) for 10 min at room temperature, then washed extensively with distilled water.

### Hyaluronidase activity measurement

Hyaluronidase activity was measured as described previously (76, 112). Briefly, pneumococcal cultures were grown in Columbia broth to an absorbance of 0.7 at 600 nm. Then the bacterial pellet and supernatant were obtained by centrifugation. Prior to the reaction, bacterial pellets were washed thoroughly with PBS at room temperature then diluted in 250 μl of assay buffer while supernatant samples were diluted 1:1 in assay buffer. Hyaluronic acid isolated from *Streptococcus equi* (Sigma Aldrich) was utilized at a final concentration of 10 ug ml^-1^ in assay buffer (150 mM NaCl, 200 mM sodium acetate, pH 6) for 15 min at 37°C. Stop buffer (2% NaOH, 2.5% cetrimide) was added to stop the reaction, and the absorbance at 400 nm was determined. Commercial hyaluronidase from bovine testes (Sigma-Aldrich) was used as positive control. Assay results were calculated using the standard curve from the positive controls and by comparison to that of a blank comprised of assay buffer solely or without bacterial material.

### Mammalian cells and media growth conditions

Epithelial lung human carcinoma A549 and fibroblast NIH-3T3 cells were propagated and maintained at 37°C and supplemented of 5% carbon dioxide at pH 7 in DMEM media contained 1% L-glutamine, 5% fetal bovine serum (FBS) without antibiotics. Chinchilla middle ear CMEE cells were maintained as previously reported (7, 71).

### siRNA knockdown

All transfection reagents were provided by Sigma-Aldrich. Subconfluent A549 cultures (20% to 30%) were seed in 24-wells plates and transfected with 3 pmol of a mixture containing pre-designed siRNA. Each mixture consisted of multiple siRNA targeting specifically either *HAS1*, *HAS2*, and *HAS3*. Universal control siRNA was used as a negative control. siRNA was transfected in a final volume of 100 μl of OptiMEM containing 3 μl of MISSION siRNA transfection reagent and no antibiotics. Twenty-four hours later the medium was replaced by DMEM supplemented with fetal bovine serum (FBS). Forty-eight hours after transfection, cells were used in adherence assays. To verify the reduction in *HAS* expression, A549 cells were lysed using the RNeasy kit and relative *HAS1*, *HAS2*, *HAS3*, and *ACTINB* expression was verified by qPCR, as described above.

### Pneumococcal host cell adherence assay

A549, NIH-3T3 or CMEE were seed in 24-wells plates or on Lab-Tek II Chamber Slides (Thermo Fisher) at a density of 1 x 10^5^, and maintained without antibiotic until confluence. Before infection, the cell monolayer medium is replaced with fresh medium and then infected with pneumococci resuspended in DMEM supplemented with 1% FBS using a multiplicity of infection (MOI) of 50 bacteria per cell. Infected cells were incubated at 37°C in 5% CO_2_ for 1 h. Then the media of post-infected cells was aspirated and washed thoroughly three times with PBS to remove unbound bacteria. The total number of adherent bacteria was released with saponin (1% w/v) for 15 minutes at 37 °C and plated on TSA plates followed by colony formation and enumeration. The effect of exogenous HA on attachment interference was performed through the addition of HA from *Streptococcus equi* at concentrations ranging between 0.01 to 1 mg/ml to the A549 monolayers before the addition of the bacteria. Each experiment was repeated at least four times, and results were expressed as mean ± S.E.M.

### Confocal microscopy

A549 cells were seeded on Nunc Lab-Tek 4-chamber (VWR) and inoculated with pneumococcal strains with a MOI of 50 as described above. After 1 hour of infection, cells were washed 3-times with PBS and fixed with 4% PFA, washed with PBS and stained with BODIPY FL Vancomycin (Thermo Fisher Scientific), Alexa Fluor 568-Phallodin (Thermo Fisher Scientific) following manufacturer’s instructions. Finally, cells were covered with Vectashield mounting media with DAPI (Vector Laboratories). Confocal microscopy was performed on the stage of Carl Zeiss LSM-880 META FCS, using 488 nm laser line for the fluorescence tag and 561 nm laser line for the fluorescence Alexa tag. Images were analyzed using ZEN Black software (Carl Zeiss Microscopy, Thornwood, NY) and assembled in Fiji (ImageJ. 2.0.0rc48). Laser intensity and gain were kept the same for all stacks and images.

### In vivo studies

Colonization ability of pneumococcal strains were tested essentially as previously described (83, 84). Nine-week-old CD1 mice were raised in the pre-clinical facility of the Leicester University. Mice were anesthetized lightly with 2.5% isoflurane over oxygen (1.5 to 2 liter/min), and they were inoculated intranasally with 20 μl PBS, pH 7.0, containing approximately 5X10^5^ CFU/mouse. When inoculating, mice were kept horizontally to prevent the dissemination of inoculum into the lower respiratory tract. A group of mice (n = 5) for each cohort were killed by cervical dislocation at 2 hours and 7 days after infection. After removing the mandible, nasopharyngeal cavity was accessed using a 18G needle through the soft palate, and the content of nasopharynx was collected by injecting 500 μl PBS, pH 7.0. This process was repeated once more using the same lavage fluid in order to optimize the recovery of bacteria. Undiluted and serially diluted lavage fluid was plated on blood agar base plates and CFU/nasopharynx was calculated. The results were analyzed by two-way ANOVA followed by Tukey’s multiple comparisons test.

### Ethics statement

Mouse experiments were done under the UK Home Office approved project (permit no: P7B01C07A) and personal (permit no. I66A9D84D) licenses. The protocols used were approved by both the UK Home Office and the University of Leicester ethics committee. We used anesthetic during the procedure to minimize the suffering. Moreover, the animals were kept in individually ventilated cages, had access to water and feed *ad libido*, and their health were checked regularly in accordance with the legal and institutional requirements.

### Statistical analysis

Statistical significance was assessed using Student’s two-tailed *t*-test for independent samples, one-way ANOVA and Bonferroni’s multiple comparisons test for independent samples, and two-way ANOVA and Tukey’s multiple comparisons test for animal data. *p*-values less than 0.05 were considered statistically significant.

## Acknowledgments

We are very grateful to Dr. Aaron Mitchell and Dr. Frederick Lanni for their insightful suggestions and guidance during this project. We also thank Dr. Samantha King for productive discussions, particularly in regards to hyaluronic acid. We thank Dr. Haibing Teng from the Molecular Biosensor and Imaging Facility (MBIC) at Carnegie Mellon University for her assistance with confocal imaging. We thank Dr. Phil G. Campbell for providing A549 cells. This work was supported by NIH grant R01 AI139077-01A1 to N.L.H., the Eberly Family Career Development Professorship of Biological Science, an award from the Samuel and Emma Winters Foundation, as well as support from Carnegie Mellon University. The authors declare that there are no conflicts of interest.

## Author Contributions

RAC contributed to the conception, design, experimentation, analysis and interpretation, and manuscript preparation; EE contributed with design and construction of bacterial clones, as well as data interpretation; OG contributed with *in vivo* experiments and data analysis, HY contributed with *in vivo* experiments, analysis and interpretation, and manuscript preparation; NLH contributed design, analysis and interpretation, and manuscript preparation.

---

**Fig S1.**
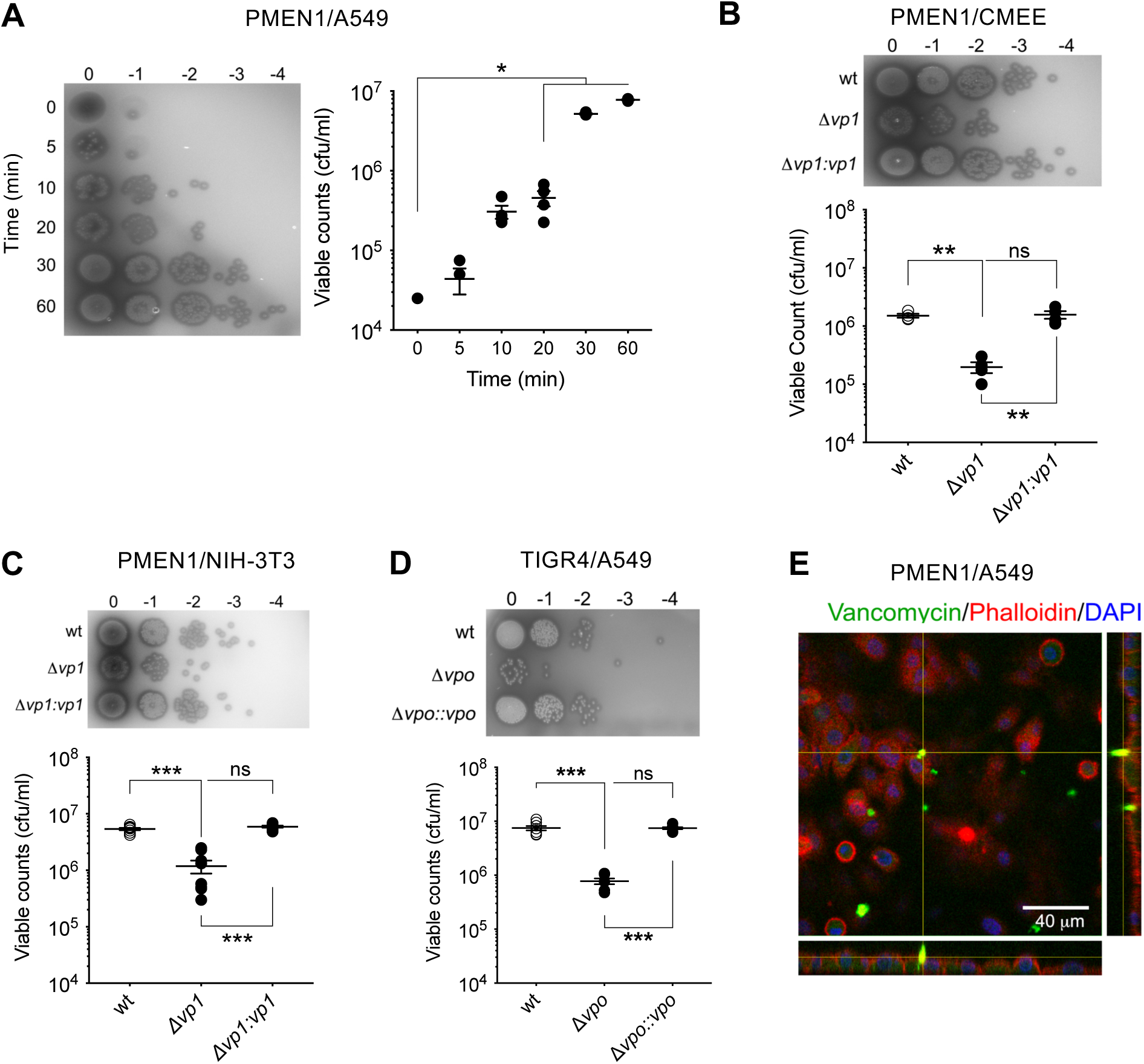
*vp1* enhances pneumococcal attachment to epithelial cells. (A) A549 cells were inoculated with the wild-type PMEN1 strain PN4595-T23. After washing the unbound bacteria, the total cell-bound bacteria were recovered. For visualization, five microliters of ten-fold dilutions were used for spot assays on TSA plates (left panel). For quantification, the total number of bacteria was determined by enumeration on TSA plates (right panel). (B and C). Attachment assay were performed on CMEE cells (B) and NIH-3T3 cells (C) inoculated with the strain wild-type PN4595-T23, the *vp1* mutant and *vp1* overexpressor strain (Δ*vp1* and the Δ*vp1:vp1,* respectively). As in A, cell-bound bacteria were recovered and enumerated on TSA plates (bottom panels). (D) A549 cells were inoculated with the wild-type TIGR4 strain and its derivative *vpo* mutant and *vpo* overexpressor strains (Δ*vpo* and the Δ*vpo::vpo,* respectively). The total cell-bound bacteria were recovered and enumerated as in A. (E) Representative orthogonal view image of A549 cells inoculated with wild-type PN4595-T23. Note the extracellular *foci* indicating surface binding of the bacteria. Bacterial cells were stained with BODYPI-FL vancomycin (green), actin was visualized with TRITC-phalloidin (red), and bacterial and human DNA was visualized with DAPI (blue). Data on A, B, C, and D represents the mean ± S.E.M of at least four independent experiments. Statistical significance was calculated using unpaired one-way ANOVA analysis with Bonferroni correction, and multiple comparison between samples were performed. * *p* < 0.005, ** *p* < 0.001, *** *p* < 0.0001, ns = not significant.

**Fig S2.**
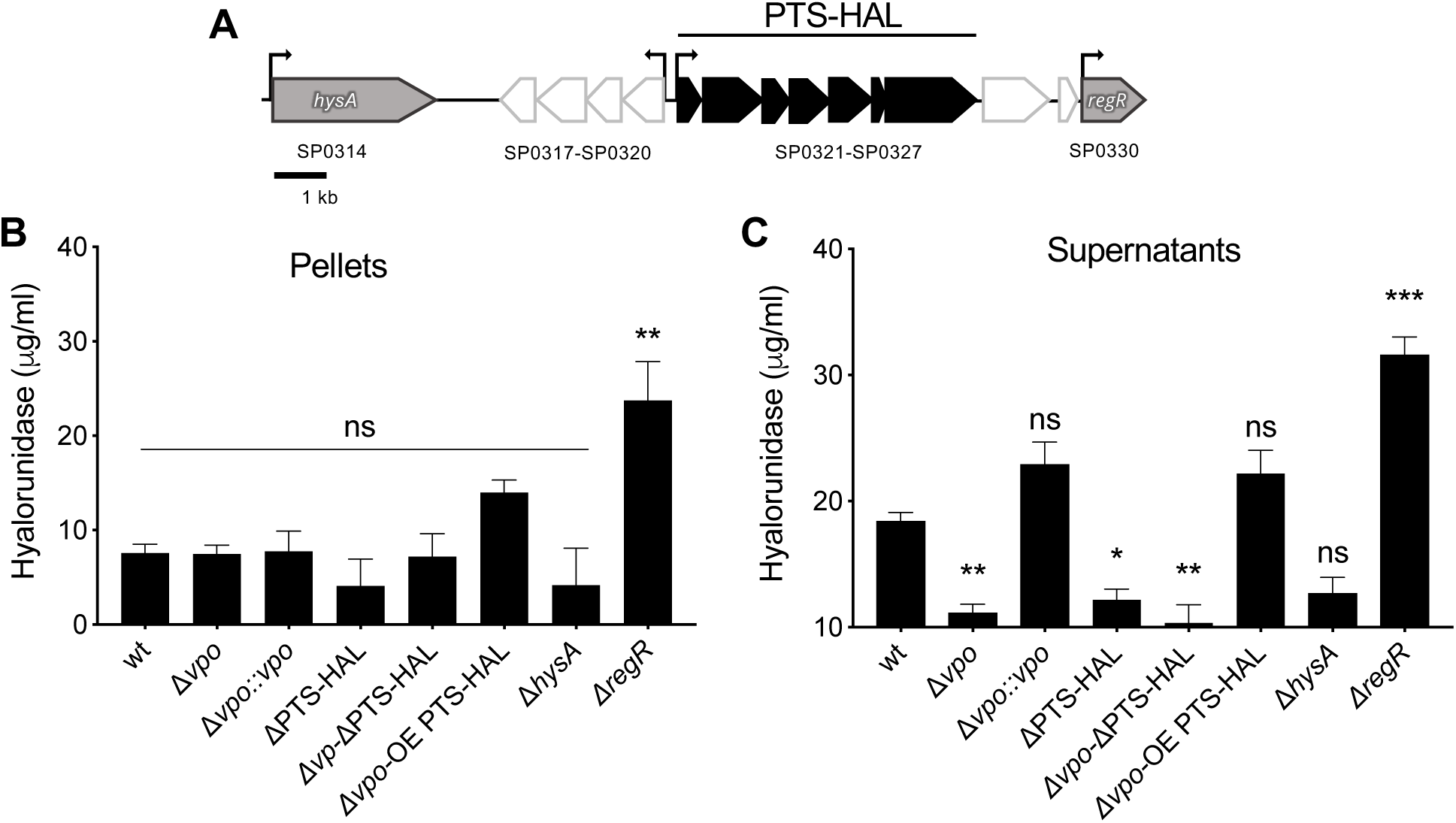
The *vp1* product promotes degradation of the hyaluronan polymer via its controls of the operon encoding the PTS-EII system in TIGR4. (A) Schematic of the genomic region implicated in the metabolism of hyaluronic acid. Arrows correspond to predicted open reading frames in TIGR4. Black and white denote predicted operons; PTS-HAL has two additional open reading frames at the end of the predicted operon (represented in white). Grey corresponds to genes involved in HA metabolism, but not highly controlled by *vp1*. The IDs refer to gene names in the TIGR4 genome. (B and C) Measurements of breakdown of hyaluronan by strain TIGR4 and isogenic mutants. Both the supernatants (B) and the pellets (C) of bacteria grown in liquid cultures were collected and assessed for hyaluronidase activity relative to the wild-type strain. Data on B and C represents the mean ± S.E.M of 8 independent experiments. Statistical significance was calculated using unpaired one-way ANOVA analysis with Bonferroni correction, and multiple comparison between samples were performed. * *p* < 0.05, ** *p* < 0.05, *** *p* < 0.0001 ns = not significant relative to the wild-type strain. Note that on C, the Δ*vpo:vpo* rescues the hyaluronidase deficiency of the Δ*vpo* mutant (*p* < 0.0001) and the overexpressor Δ*vpo-*OE PTS-HAL rescues the hyaluronidase activity of the mutants Δ*vpo* (*p* < 0.0001). Further, Δ*vpo-*OE PTS-HAL is also significantly different from the double-mutant Δ*vpo-*ΔPTS-HAL (*p* < 0.0001).

**Fig S3.**
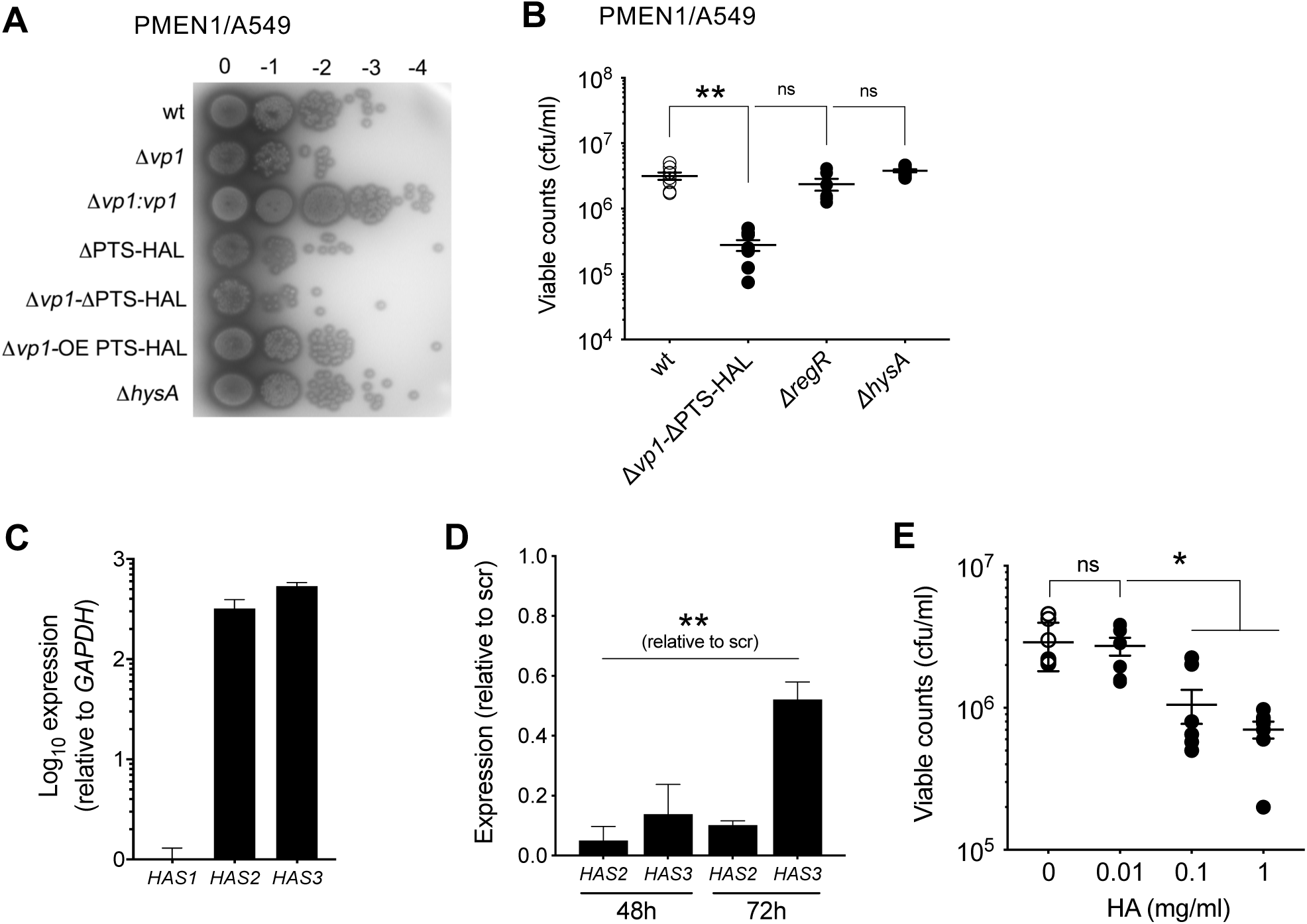
Pneumococcus attaches to host HA in a process controlled by VP1 and mediated by molecules involved in hyaluronic acid processing. (A) Spot assay to access attachment to host cells. The wild-type PN4595-T23, and its derivative strains Δ*vp1,* Δ*vp1*:*vp1,* ΔPTS-HAL, Δ*vp1*-ΔPTS-HAL, Δ*vp1*-OE PTS and Δ*hysA*, were assessed for attachment to A549 cells using a spot assay. After washing the unbound bacteria, the total cell-bound bacteria were recovered and plated on TSA plates. (B) Quantification of binding, where the total number of the Δ*vp1*-ΔPTS, Δ*regR and* Δ*hysA,* bound to epithelial cells was enumerated on TSA plates. (C) Measurement of the expression levels of hyaluronic acid synthases-encoding genes *HAS1*, *HAS2*, and *HAS3* in A549 cells at two time points where levels are compared to treatment with a scrambled control. (D) Measurement of the expression levels of hyaluronic acid synthases-encoding genes *HAS2* and *HAS3* after treatment with specific siRNA. (E) Quantification of bacterial binding to A549 pre-incubated with varying concentration of HA. Data on B to E represents the mean ± S.E.M of at least four independent experiments. Statistical significance was calculated using unpaired one-way ANOVA (B, D, and E) analysis with Bonferroni correction, and multiple comparison between samples were performed. * *p* < 0.005, ** *p* < 0.0001, ns = not significant.

**Fig S4.**
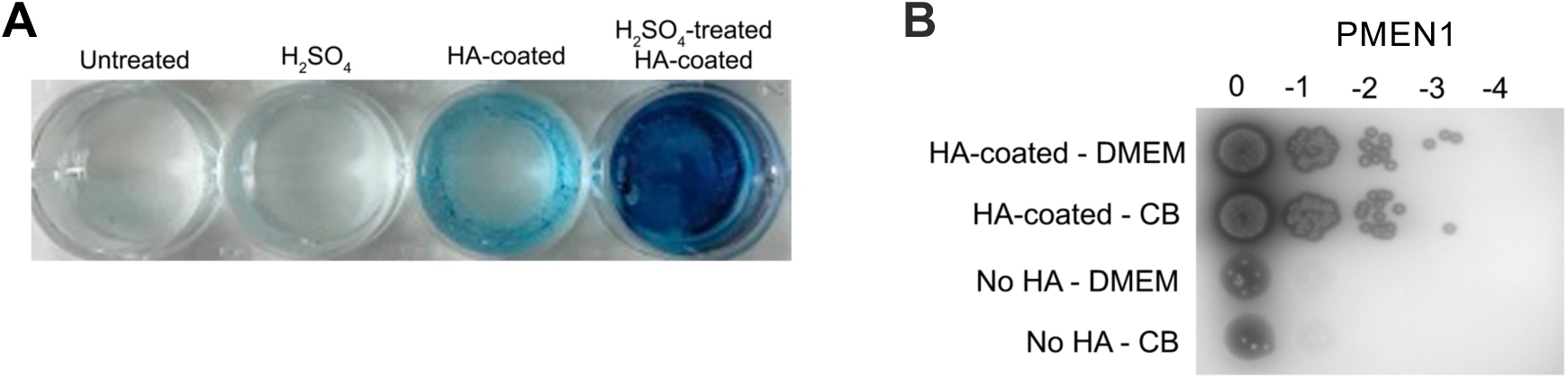
*vp1* mediates attachment to hyaluronic acid of *Streptococcus equis* via genes in the PTS-HAL operon. (A) hyaluronic acid (HA) coating of 6 well plates under listed treatments, visualized by staining with Alcian Blue 8GX. (B) The PMEN1 strain PN4595-T23 (top panel) was assessed for attachment to hyaluronic acid-coated surfaces in FBS-supplemented DMEM media (A549 growth media) or Columbia broth (bacterial growth media). The total cell-bound bacteria were recovered and tested for growth on TSA plates.

